# Conformational State of Myosin’s Disordered Loop 2 Structure Mediates Actomyosin Association During Crossbridge Formation

**DOI:** 10.1101/2025.08.13.670102

**Authors:** Kalen Z. Robeson, Matthew Carter Childers, Kieran J. Fruebis, Rachelle Soriano, Jennifer Davis, Michael Regnier

## Abstract

The binding of myosin to actin to form crossbridges is a critical step for force generation by sarcomeres. A recent cryo-electron microscopy structure has resolved the weakly-bound actomyosin complex (AM.ADP.Pi); however, the structural and dynamic factors that influence actin-myosin association are unclear. The disordered loop 2 of myosin is thought to mediate actomyosin interactions in complex, chemomechanical state-dependent fashion; however, the loop is usually unresolved in structural studies due to its intrinsic disorder. Here, we utilize a combination of molecular dynamics simulations and electrostatic calculations to investigate the dynamics of these actin binding regions of myosin. Our results show that loop 2 experiences disordered dynamics and that specific conformations sampled by loop 2 modulate the strength of the associative electrostatic force between actin and myosin. Variation in the actin-myosin associative force was associated with the presentation and orientation of positively charged residues in loop 2. We provide an in-depth analysis of the conformational state space occupied by loop 2 during nine 500 ns molecular dynamics simulations of pre-powerstroke human β-myosin S1, with three replicates each of wildtype and two different mutant (E525K and V606M) myosin structures. This dataset allowed for exploration of how loop 2 conformational sampling is altered by these two mutations which have been clinically and experimentally associated with cardiomyopathy and altered actin binding affinity. The E525K and V606M mutations altered the conformational ensemble sampled by loop 2 and were associated with associative actin binding strength. These results highlight the importance of the positive charges on loop 2 for actomyosin interactions and demonstrate how disease-causing mutations outside of loop 2 can still affect it.

## Introduction

Contractions in muscle are a product of myosins binding to actin to form crossbridges that cycle to produce force and shorten. In cardiac muscle, identifying the residues in myosin that interact with actin before and during the formation of the weakly-bound actomyosin complex is a vital step towards understanding the regulation of force development and how this may be altered by disease-causing mutations, as well as for the development of novel treatments for cardiomyopathies. In the weakly-bound state (AM.ADP.Pi), the actin binding surface of myosin is made up of the helix loop helix (HLH) motif of the lower 50 kDa domain, loop 3, which is also on in the lower 50 kDa domain, and loop 2, which spans the cleft between the upper and lower 50 kDa domains (**Figure 1A**)^1,2^. Loop 4 and the cardiomyopathy loop also contribute to actin interactions in force-bearing states of the crossbridge cycle. The positively charged lysine residues on loop 2 are known to interact with negatively charged residues on the N-terminus and Asp-containing N-terminal hairpin loop of actin during weak and strong (AM.ADP*) binding ^3–7^. However, to date, no structures resolve the complete loop 2 structure for any myosin isoform or in any chemomechanical state. In this study we explore the dynamics of loop 2 in the pre-powerstroke (M.ADP.Pi) state and report on how disordered dynamics modulate the presentation of positive charges by myosin.

**Figure 1:**
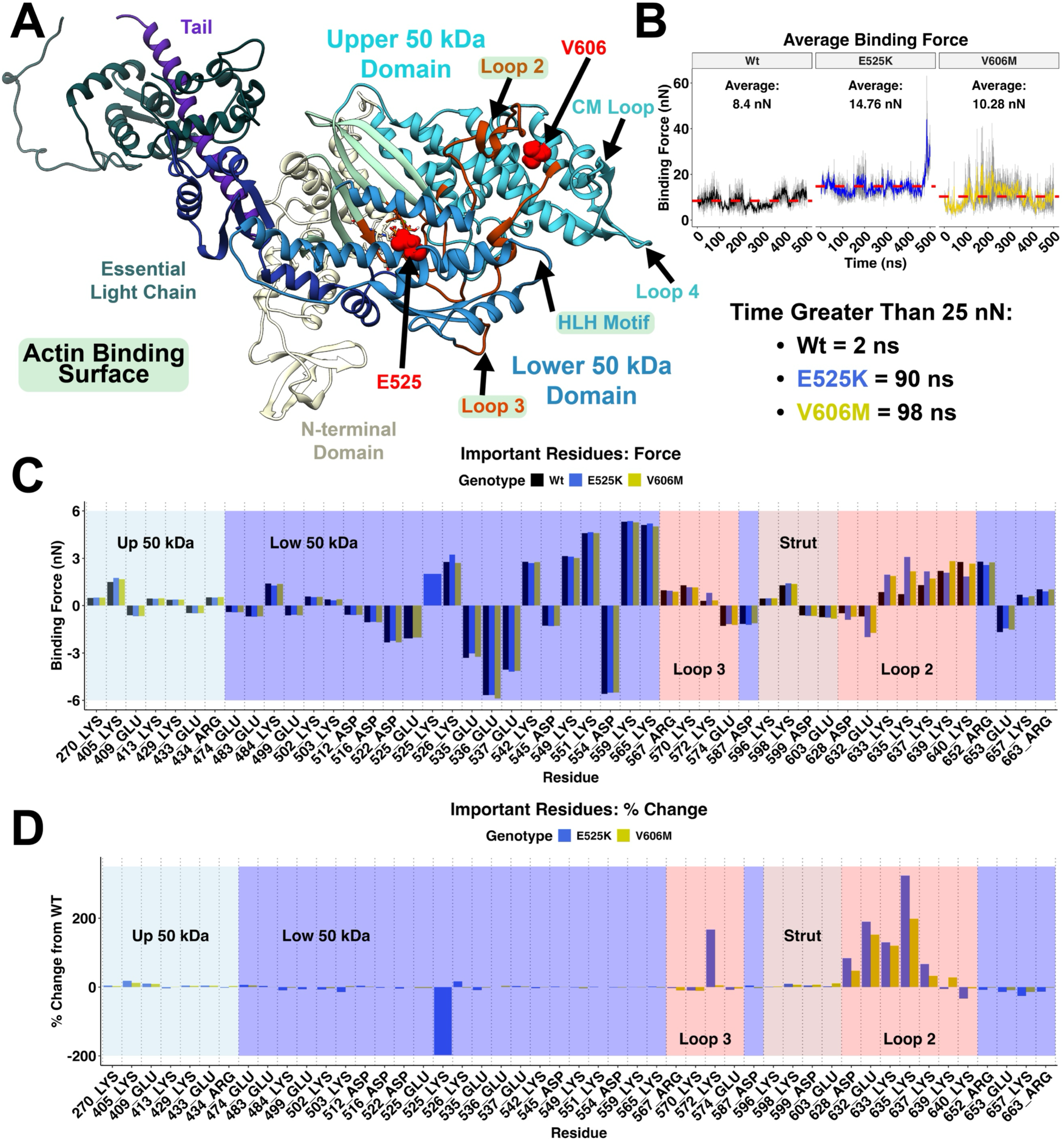
Loop 2 Drives Observed Changes in E525K and V606M Myosins’ Affinity for Actin. (A) Pre-powerstroke human β-myosin S1. The major structural domains as well as the location of the E525K and V606M mutations are labeled. Structures of the actin binding surface are highlighted in teal. (B) Average binding force calculated with *DelPhiForce* across simulation time in three replicate simulations of each genotype (Mean + SEM; dashed lines represent mean across all simulations of a given genotype. High binding energy (pro-actin binding) conformations are sampled in E525K and V606M simulations (bottom). (C) Quantification of binding force felt at the single amino acid level in residues that experienced > ±0.0 kT/Å during any (WT or Mutant) *DelPhiForce* simulation. (D) Percent change from WT binding force for each residue in C. The positively charged lysine residues in loop 2 can be seen driving increased actin affinity in E525K and V606M. The charge change due to the E525K mutation is also identified in E525K simulations.

To investigate the dynamics of the actin interacting myosin domains we developed an *in-silico* assay for actomyosin crossbridge formation by coupling molecular dynamics (MD) simulations with electrostatic calculations using *DelPhiForce*^8^. Two cardiomyopathy associated mutations (E525K and V606M) that are near, but not in, loop 2 and increase myosin’s affinity for actin^9–11^ were used to perturb myosin’s structure. Comparison of WT and mutant behavior highlighted the importance of loop 2 and led us to an in-depth structure-function analysis that explores the relationship between sequence and the conformational state space occupied by loop 2.

The two mutations surround loop 2 (residues 621-646): E525K is in the lower 50 kDa domain and V606M in the upper 50 kDa domain (**Figure 1A**). The E525K mutation involves a charge inversion, disrupting key salt bridges that connect the E525 residue to the helix where loop 2 is anchored in the lower 50 kDa domain. This mutation could therefore lead to allosteric changes in the conformational ensemble sampled by loop 2. The V606M mutation extends the hydrophobic side chain of V606 and increases the likelihood of hydrophobic interactions inside the upper 50 kDa domain. This mutation is found on a helix in the upper 50 kDa domain which leads to loop 2 and, again, could lead to allosteric changes in the conformation of loop 2 (**Figure 1A**). These two mutants perturb the structure of myosin in the region where loop 2 interfaces with the upper and lower 40 kDa domains. We hypothesize that both mutations change the dynamic presentation of the charges in loop 2.

Previous work has established the importance of loop 2 via both mutation screening^4,6^ and MD simulations of myosin^7,12^. Indeed, myosin mutations that neutralize a positively charged residues in loop 2 are associated with dilated cardiomyopathy and a decrease in myosin’s affinity for actin^13^. Despite the importance of loop 2 for crossbridge formation, this study represents the first in-depth exploration of both loop 2’s dynamic movement and the impact of that movement on electrostatic interactions between myosin and actin. We modeled the pre-powerstroke (M.ADP.Pi) conformation of human MYH7 performing three replicate simulations of each myosin variant (WT, E525K, V606M), to resolve an ensemble of loop 2 conformations. The *DelPhiForce* tool generated by Li *et al.*^8^ was then used to calculate the electrostatic force from actin on myosin with singe atom resolution. In depth analysis of these results suggested that loop 2 conformation is responsible for the changed actin affinity observed with these myosin mutations. This work provides a mechanistic groundwork for understanding the role of loop 2 and how mutations that influence its structure and mobility change myosin’s affinity for actin.

## Results

### Electrostatic Modeling Identifies Residues That Initiate Actin Binding and Highlights the Importances of Loop 2 in Crossbridge Formation and Cardiomyopathy

First, we performed conventional MD simulations of WT, E525K, and V606M human cardiac β-myosin in the pre-powerstroke (M.ADP.Pi) state. The *DelPhiForce* tool developed by Li *et al.* was used to calculate the electrostatic forces that occurred between a model actin filament (an unregulated pentamer) and different MD-derived conformations of myosin^8^. Structures from MD simulations of pre-powerstroke human β-myosin S1 were sampled at 1 ns resolution (500 structures per simulation) and aligned 30 Å away from the central myosin binding sight on the actin pentamer (**Supplemental Figure 3A**). *DelPhiForce* was then used to calculate the electrostatic attraction felt by myosin from actin for each atom in the myosin model for each sampled snapshot.

The calculated binding force for each timestep in replicate simulations for both WT and mutant myosins was variable (**Figure 1B and Supplemental Figure 1B**), but the average binding force calculated across all E525K and V606M timesteps and simulations replicates was elevated compared to WT (**Figure 1B** dashed lines). Additionally, high binding energy (>25 nN) conformations were sampled more frequently in E525K and V606M simulations compared with WT simulations (**Figure 1B**). To investigate the electrostatic contribution of each residue to binding force calculations, residues that resulted in > ± 0.01 Kt/Å binding energy during any *DelPhiForce* calculation were selected for further analysis (**Supplemental Figure 1A**). The analysis allowed us to define residues that have electrostatic interactions with actin during early myosin binding to actin with high resolution. We identified charged residues in known actin binding elements including the HLH motif, loop 2, and loop 3. Quantification of the binding force from each of these residues shows that the positively charged residues in the HLH motif and loop 2 experienced the strongest attractive force (**Figure 1C**). To predict the effects of E525K and V606M on myosin affinity for actin, we calculated the precent change in electrostatic force between the WT and mutant simulations. For most residues, the attractive or repulsive force on myosin from actin was relatively unchanged between WT and mutant simulations. However, the positively charged Lys residues in loop 2 had the greatest changes in binding force compared to WT (**Figure 1D** and **Supplemental Figure 1**). Also, as expected, the charge inversion of the E525K mutation led to a shift from a repulsive force to an attractive force at this location (**Figure 1D**).

Together these results indicate that isolated, static structures are not adequate to explain the changed actin affinity seen with these mutations. Instead, conformational ensembles are required to capture the range of electrostatic forces between actin and myosin during the early stages of crossbridge formation. The temporal and spatial resolution provided by MD simulations more accurately samples the relationship between structural heterogeneity and actomyosin interaction of loop 2. The changes in actin affinity appear to be driven by conformations of loop 2 that are only seen in E525K and V606M simulations and never sampled by WT simulations. This result prompted a deeper investigation of how these mutations alter loop 2 movement and how this relates to myosin activation and crossbridge formation.

### The Dynamics of Loop 2 are Changed by The E525K and V606M Mutations

We next investigated the hypothesis that myosin mutations near the interface of loop 2 with the upper and lower 50 kDa domains can disrupt its movement and the presentation of loop 2’s positive charges to actin. Contact pair analysis of E525K and V606M MD simulations showed that the contacts formed by loop 2 during simulations were changed by both mutations. The E525K mutation in the lower 50 kDa domain resulted in a charge change (negative to positive) and disrupted salt bridges on adjacent helices (**Figure 2A-B**). One of these helices leads directly to loop 2, and the contacts formed by loop 2 during MD simulations of E525K myosin were significantly altered compared to WT simulations (**Figure 2C**). In MD simulations of V606M myosin the increased hydrophobic side chain length induced a hydrophobic interaction between the 606 reside and residue 434 on an adjacent helix in the upper 50 kDa domain (**Figure 2D-E**). This interaction twisted the helix where residue 606 is located and led to allosteric shifts in loop 2 as seen by altered loop 2 contacts in V606M simulations (**Figure 2F**). This analysis showed that there are intramolecular pathways where each of these mutations changes the allostery of and contacts formed by loop 2.

**Figure 2:**
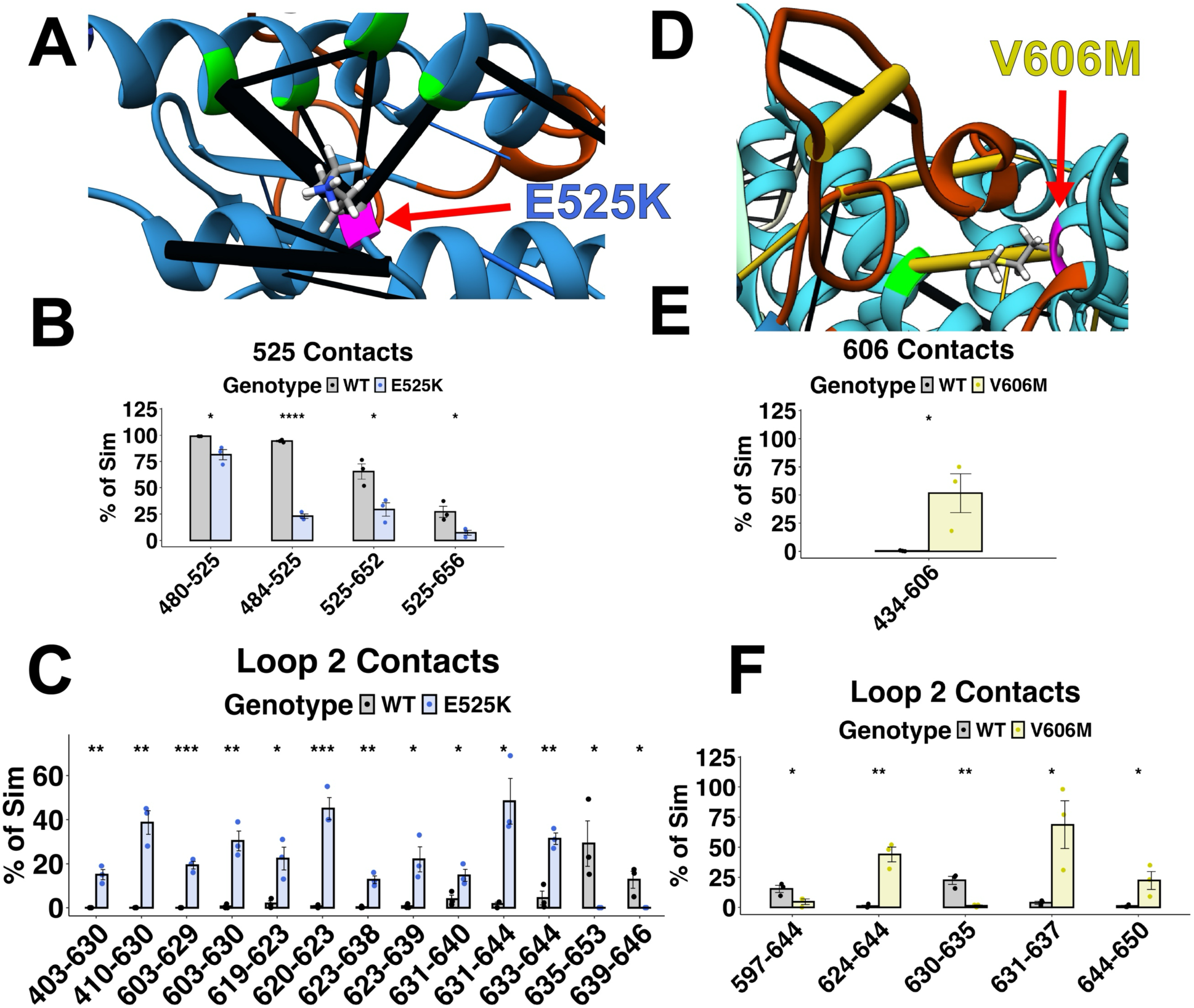
The allostery of myosin’s loop 2 is disrupted by the E525K and V606M mutations. (A) Zoomed in contact pair image of residue 525 in myosin showing how the E525K charge change disrupts key salt bridges on adjacent helices. Blue bars indicate increased contact time and black bars indicate decreased contact time compared to WT. (B) Quantification of contact pairs involving residue 525 during the WT (black) and E525K (blue) simulations. (C) Quantification of altered contact pairs involving loop 2 during E525K simulations compared to WT simulations. (D) Zoomed in in contact pair image of residue 606 showing how the V606M mutation forms a hydrophobic interaction with residue 434. Yellow bars indicate increased contact time and black bars indicate decreased contact time compared to WT. (E) Quantification of contact pairs involving residue 606 during the WT (black) and V606M (yellow) simulations. (F) Quantification of changed loop 2 contact pairs during V606M simulations compared to WT simulations. Contact pairs that were present for significantly (p < 0.05, two-tailed Student’s t-test) more or less time than WT simulations are depicted. The pseudo-bond radii are proportional to the difference in contact time: thicker pseudo-bonds correspond to greater contact populations.

These two mutations represent a framework to study the role of not only the positive charges in loop 2 but the dynamic movement of these charges and the conformational states populated by the motion and coiling of loop 2. We hypothesize that the conformational ensemble sampled by loop 2 plays a critical role in regulating actomyosin complex formation and that E525K and V606M both alter loop sampling. In turn these changes to Loop 2 conformation may result in changed electrostatic interaction between myosin and actin observed in both mutations.

### The E525K and V606M Mutations Shift Loop 2 Towards the Actin Binding Cleft

Loop 2 sampled many different conformational states during the 500 ns simulations. To identify and compare characteristic conformations (representative states) observed for WT and mutant myosin we performed principal component analysis (PCA), focusing on the Cartesian coordinates of the C_α_ atoms for each amino acid within loop 2. This dimensionality reduction allowed us to group similar structures along the different principal components. Principal component 1 (PC1) explained 45% of the observed variance and separated most mutant simulations from WT simulations, except for one V606M simulation. PC2 and PC3 did not show separation of WT and mutant simulations but described distinct types of loop 2 movement (**Figure 3A**). The contribution of each residue’s Cartesian coordinate to each principal component helps to explain the dynamic way in which these two mutations changed the conformational state space occupied by loop 2. PC1 was dominated by a negative shift in the x-coordinate of loop 2 C_α_ positions. This represents a movement of loop 2 parallel to the HLH motif and actin binding surface (**Figure 3B**). Since this principal component separates most mutant from WT simulations it appears that both mutations led to loop 2 spending more time near the tip of the 50 kDa domain and the actin binding cleft. PC2 is dominated by a positive shift in the y-coordinate of the loop 2 C_α_ position associated with movement that was orthogonal to the HLH motif and actin binding surface (**Figure 3C**). Notably V606M simulations separated from the WT and E525 structures along this axis. Finally, PC3 is made up of a negative shift in some loop 2 C_α_ coordinates and a positive shift in other loop 2 C_α_ coordinates describes a twisting of the flexible loop 2 structure (**Figure 3D**).

**Figure 3:**
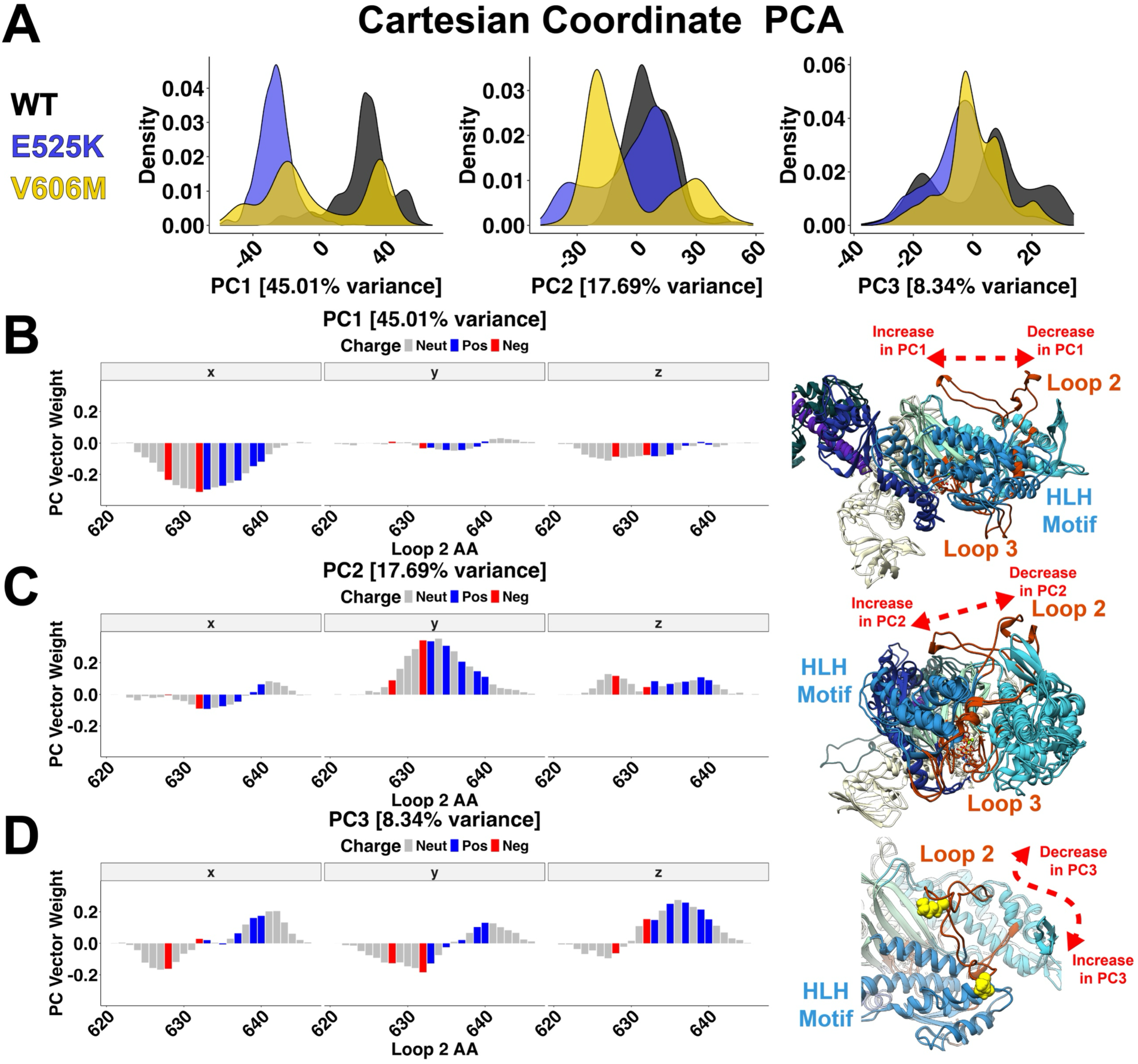
Movement of loop 2 is shifted towards the actin binding cleft by the E525K and V606M mutations. (A) Principal component analysis of loop 2 C_α_ position during 3 replicate simulations of each genotype showing mutant vs. WT separation primarily along PC1 with some contributions from PC2 and PC3. (B-D) Principal component weight analysis (left) with representative (min and max PC weight) structures from one WT simulation (right). PC1 (B) describes waving of loop 2 in the x direction (left) representing movement along the myosin head (right). PC2 (C) describes waving of loop 2 in the y direction (left) representing movement up and down between the 50 kDa domains (right). PC3 (D) describes twisting of loop 2 in x, y, and z (left) describing twisting of loop 2 (right).

These results demonstrate a method for quantifying the movement of loop 2 through time during MD simulations and show how two different myosin mutations, on either side of loop 2, can have similar allosteric impacts on its movement. They also suggest how these allosteric changes in loop 2 impact the presentation of loop 2’s positive charges, and may be important for actomyosin complex formation. As such, this result promoted a much deeper analysis of how loop 2 coils and twists during simulations of pre-powerstroke myosin.

### Expanded Conformations of Loop 2 Favor Actin Binding

Thus far, simulations demonstrated that loop 2 was responsible for changed electrostatic interactions with the E525K and V606M mutations and, in turn, these mutations changed the position of loop 2 in similar ways. We next investigated how the specific conformation of loop 2 may change the presentation of positive charges in the loop. We correlated the distance between the center of mass of positive charges in loop 2 and negative charges the N-terminus of actin (spheres in **Figure 4A**) with the binding force calculated for the whole myosin S1 head (**Figure 4B**). This correlation showed that the position of loop 2 alone explains 62% of the calculated binding energy for the whole myosin S1 head. Additionally, single timesteps from E525K simulation 2 and V606M simulation 3 can be seen that occupy conformations with very low distances and very high forces.

**Figure 4:**
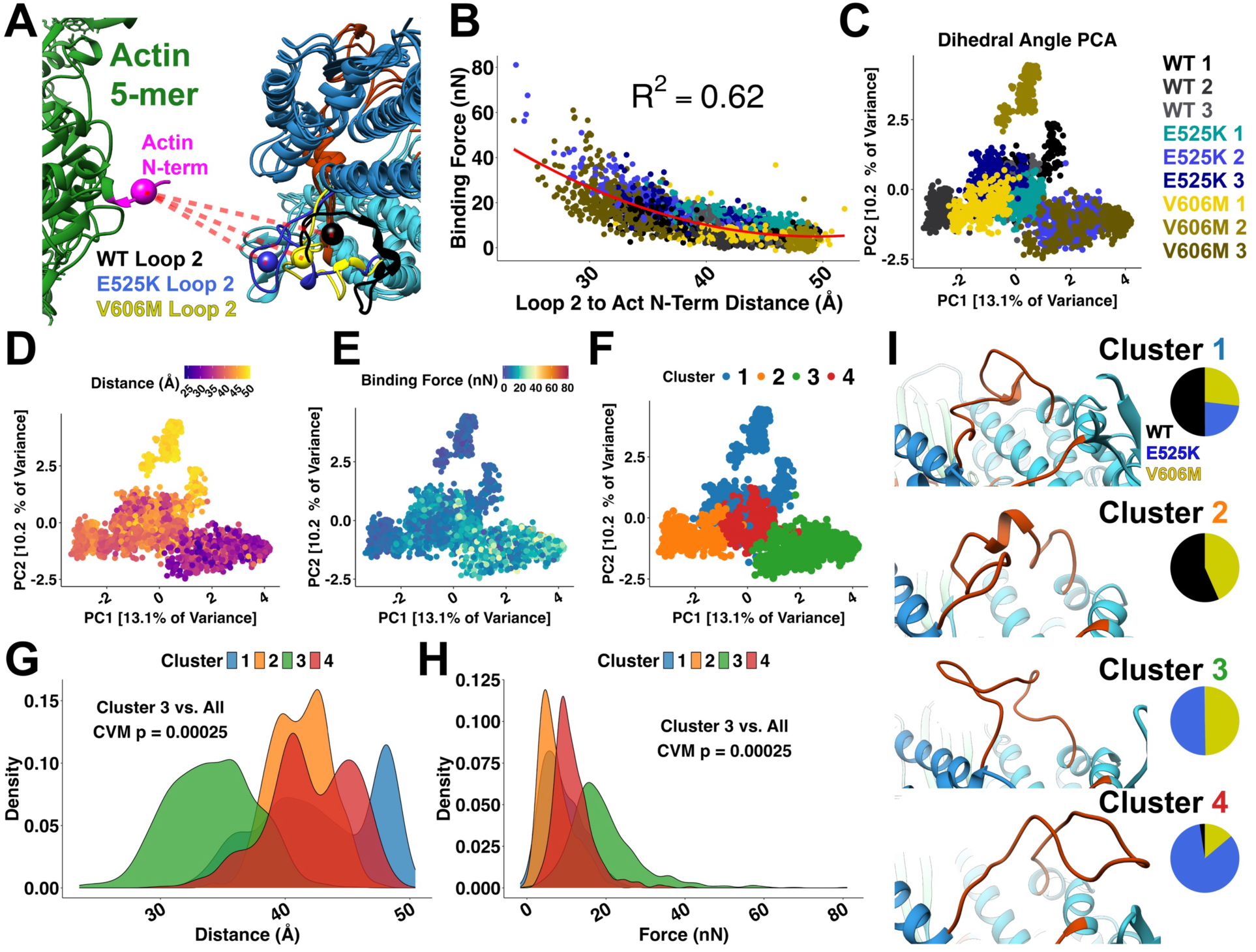
Dihedral angle analysis identifies loop 2 condensed and expanded states which regulate electrostatic binding energy. (A) Alignment of myosin 30 Å from its binding site on actin used for *DelPhiForce* electrostatic calculations. The center of mass is indicated for the negative charges on the actin N-terminus (pink sphere) and the positive charges in loop 2 (black-WT, blue-E525K, and yellow-V606M spheres). (B) The relationship between distance (spheres in A) and calculated force is fit with a second order exponential. (C) Principal component analysis of loop 2 dihedral angles during 3 replicate simulations of each genotype. E525K Sim 2 and V606M Sim 3 can be seen forming a dispersed cluster distal from all other simulations. (D) Distance and (E) force mapped onto the dihedral angle PCA clusters. The dispersed cluster of E525K Sim 2 and V606M Sim 3 contains structures with unusually low distances and unusually high forces. (F) K-means clustering on the dihedral angle PCA identified 4 clusters. (G) Density of distance values (spheres in A) measured within each cluster. Cluster 3 shows a significant shift towards lower distances between loop 2 and the actin n-terminus. (H) Density of force values calculated within each cluster. Cluster 3 shows a significant shift towards higher binding energy states—representing a shift to a pro-actin binding conformational state of loop 2. A Cramer Von Mises Test (CVM) was used to test for differences in the distribution of distance and force. (I) Representative structures and genotypic make up for each cluster. Representative structures were selected as those that most closely matched the mean force value calculated within each cluster.

Loop 2 is long and flexible enough that it can form both interactions with itself and interactions with other myosin domains. To compare more compact vs more extended conformations of loop 2, we performed dihedral angle principal component analysis (PCA) on the backbone (ɸ/ψ) dihedral angles along loop 2. In comparison to the C_α_ PCA discussed earlier, this analysis used an internal reference frame within loop 2’s structure, enabling a deeper analysis of internal versus overall motion in the ensembles sampled by loop 2. Results of this dihedral angle analysis showed that each simulation formed a dense cluster in PC space. Notably, E525K simulation two and V606M simulation three formed a less dense cluster near each other and away from all other simulations (**Figure 4C**). This indicates that these mutations may alter the conformational state space occupied by loop 2, populating conformations that are never or seldom seen in the WT structures.

The importance of these different conformational states can be seen by mapping the distance and force values calculated earlier onto the dihedral angle PCA plot (**Figure 4D & E**). Most simulations are tightly clustered with themselves and occupy high distance, low force conformations for most of the simulation time. However, the less dense cluster formed by E525K simulation two and V606M simulation three contains simulation timesteps with unusually small distances between loop 2 and actin’s N-terminus (**Figure 4D**) and unusually high electrostatic force (**Figure 4E**).

K-means clustering was used to define four conformational states that were occupied by loop 2 during these nine simulations (**Figure 4F and Supplemental Figure 4A**). The optimal k-value for clustering was based on the elbow point in the inertia plot of within-cluster sum of squares (**Supplemental Figure 4B**). By plotting the distance (**Figure 4G**) and force (**Figure 4H**) calculated earlier for each timestep within a cluster, we can see that cluster three stands out with significantly lower distance and significantly higher force compared to all other clusters. The mechanism behind this difference is elucidated by representative structures from each of the other clusters (**Figure 4I**). Structures from cluster one and cluster two formed helical twists where loop 2 interacted with itself. This twisting sequestered the positive charges in loop 2 away from the actin binding cleft and the negative charges present in the N-terminus of actin. Notably, timesteps from WT simulations made up the majority of these two “unfavorable actin-binding” states. The representative structure from the less dense cluster three was expanded, not twisted and shifted towards the actin binding surface and the HLH motif. This expanded state moved the positive charges in loop 2 much closer to the negative charges in actin’s N-terminal domain and represents a “favorable actin binding” state. Finally cluster four was also expanded but here loop 2 was not shifted directly toward the myosin binding surface and we observed only a modest increase in binding force for this cluster (**Figure 4H**). Importantly both expanded, “favorable actin-binding” states identified here were populated almost exclusively by mutant (E525K or V606M) simulations.

These results collectively demonstrate that conformational states of loop 2 (defined by dihedral angles) when combined with electrostatic modeling can predict myosin’s affinity for actin. These findings also underscore loop 2’s significant role in regulating actomyosin cross-bridge formation and clearly illustrate how mutations near loop 2 can allosterically influence its conformation, consequently affecting myosin’s binding affinity for actin. The ability of loop 2 to sample conformations that have variable long-range electrostatic interactions with actin may play an important regulatory role in myosin activation, and here are demonstrated to be causal in the increased actin affinity observed in two cardiomyopathy mutations.

## Discussion

### Loop 2 Electrostatically Mediates Actomyosin Interaction

Protein loops are often regulators of function and enzymatic activity in biological systems,^14,15^ and structural disorder of a loop often imparts functional benefits^16^. Spudich and coworkers established that changes in the sequence, length, and charge of loop 2 affects actin-myosin kinetics^17^ as well as ATPase activity^4,6^. While these vary substantially across myosin classes, loop 2 consistently demonstrates impact on actin-myosin association for many myosins. Yu *et al.* used AI-generated models and MD simulations of myosin to explore the relationship between loop 2 length and actomyosin interaction^18^. They report that longer loop 2 isoforms formed a more complex salt bridge network (involving actin’s N-terminus and a hairpin turn in actin subdomain 1) than did short loop 2 isoforms (few to no interactions with actin’s N-terminus) in strongly-bound states^19^. Yengo and Sweeney studied variants of the processive myosin V^20^. They observed that changing the charge on loop 2 modified the kinetics of actin-myosin binding with fewer effects on other steps of the cycle. They also postulated that the length and charge of loop 2 is essential for the processive properties of myosin V. Elfrink *et al.* studied the single-headed, processive type IX myosin that contains a >100 residue insertion in loop 2^21^. They found that deletion of the loop 2 insertion or increased ionic strength diminishes processive behavior. Prior studies from this group also show that loop 2 assists with the directionality required for processive motion in a single headed myosin^22^. Onishi *et al.* tested the molecular mechanisms by which mutations in loop 2 affect actin-activated ATPase and speculate that charged residues in loop 2 contribute to complex formation and that hydrophobic residues contribute to the coupled process of myosin cleft closure and phosphate release^23^. Thus, across multiple myosin isoforms, loop 2 is shown to have a net positive charge, a role in mediating actin-myosin association, and a substantial impact on cross-bridge cycling kinetics.

To date, few studies have explored the relationship between loop 2 structure and actin-myosin association^3,4^. This is due to the intrinsic structural disorder in the loop. Dynamic regions of proteins do not resolve well in X-ray and cryoEM structures. While loop 2 is generally unresolved in X-ray structures of myosin alone, fragments of loop 2 are frequently observed in some cryoEM structures of the actomyosin complex. Doran *et al.* solved cryoEM structures of force-bearing-like conformations of human cardiac actomyosin^24^ and their structures show residues F644-K639 interacting with the D24-D25 hairpin turn of actin. Risi *et al.* detail how changes in the length of loop 2 across myosin isoforms affects its resolution in cryoEM structures, with shorter loops being better resolved. However, N-terminal residues of actin may also not resolve in cryoEM, leaving the possibility of a ‘fuzzy cloud’ of disordered interactions^25^. In a recent study, Klebl *et al*. collected cryoEM structures of the weakly-bound, primed actomyosin complex using a myosin-V construct with mutations near the phosphate ‘back door’ and deletions in loop 2 that slow the phosphate release step^5^. Their structure shows the C-terminal end of loop 2 sandwiched between the negatively charged hairpin turn and N-terminus of actin. Together these studies provide strong evidence that loop 2 interacts electrostatically with actin in both weakly-bound and strongly bound states. However, no studies have resolved the full structure of loop 2 and none indicate the structural mechanisms through which loop 2 mediates actomyosin association. The formation of a weakly-bound crossbridge leads to stronger binding that is needed for force generation and shortening in the sarcomere as interaction between myosin and actin facilitates the lever arm swinging. The simulations performed here contain the full structure of loop 2 and our models show that loop 2 structure interconverts among multiple conformations on nanosecond timescales (**Figure 3**). *DelphiForce* measures indicate that loop 2 contributes meaningfully to the electrostatic association force between actin and myosin (**Figure 1**) and that neutralization of loop 2 charged residues diminishes this force (**Supplemental Figure 3**). Finally, our conformational analysis of loop 2 showed that loop 2 can sample compact conformations with unfavorable anti-binding properties and more extended conformations that favor actin binding (**Figure 4**). These data suggest that the conformational ensemble that loop 2 samples can regulate a key step leading to contractions by modulating initial crossbridge formation. Our results provide a model wherein the coiling of loop 2 serves as a regulatory axis for the number of crossbridges formed and force generation within the heart (**Figure 5**).

**Figure 5:**
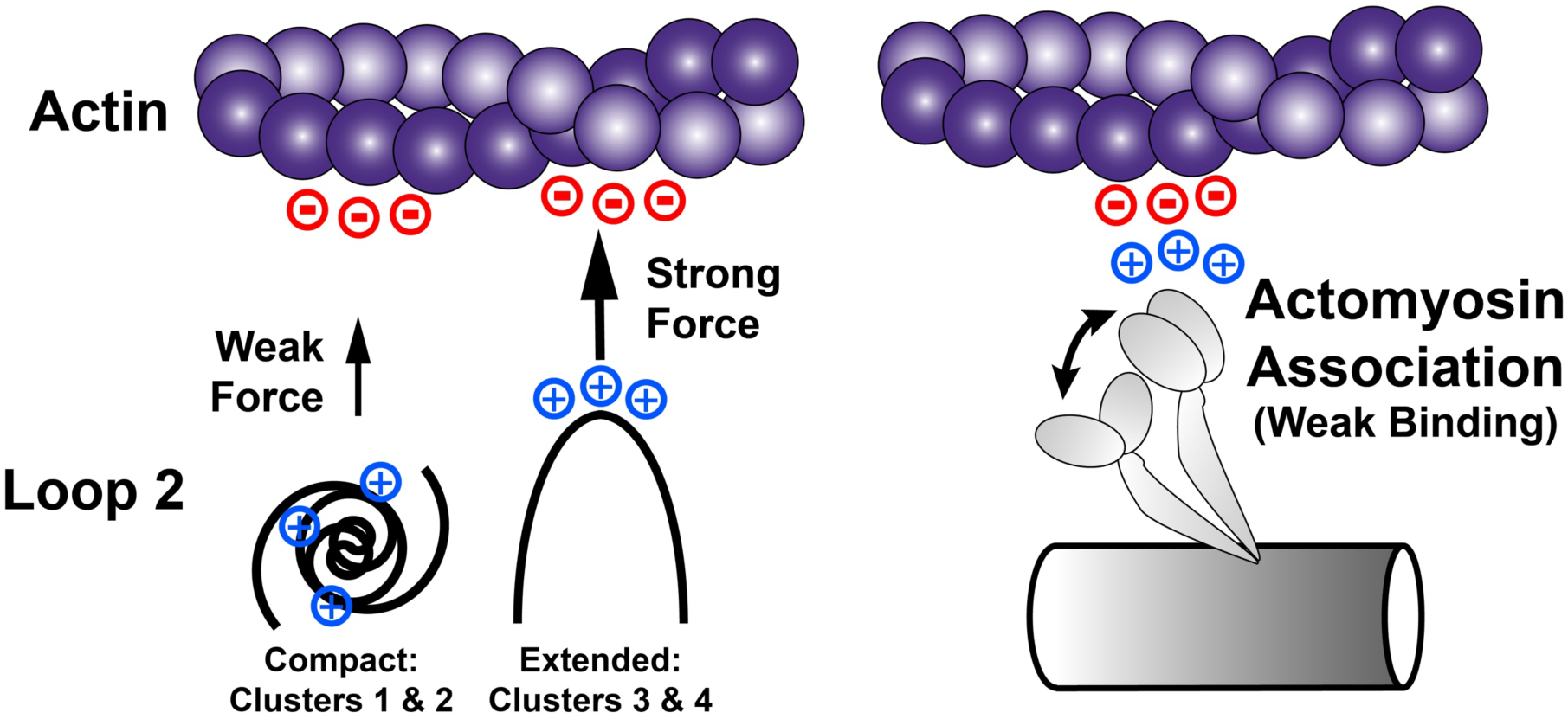
A model for the regulatory role of loop 2 in myosin activation. The loop 2 structure of myosin adopts compact semi-stable conformations where the positive charged residues in loop 2 are sequestered away from their negatively charged binding partners on the N-terminal domain of actin. Loop 2 can adopt extended less-stable conformations where these same positive changes are available to interact with actin. Here two cardiomyopathy mutations that increase myosin molecules’ affinity for actin were shown to disrupt loop 2 coiling demonstrating a mechanism for this regulatory role of loop 2 in disease progression.

### Links Between Conformational Shifts in Loop 2 and Cardiomyopathy

Over 1000 mutations in human striated muscle myosis are associated with disease^13^. This includes several mutations within loop 2 of β-cardiac myosin (e.g. K637E^26^ and K639E^27^ are associated with dilated cardiomyopathy; G636S^28^ and S642L^29^ are associated with hypertrophic cardiomyopathy). The E525K and V606M mutations studied here are not found in the loop 2 sequence and are instead located on either “side” of this loop (**Figure 1A**). However, our results show that these mutations can result in structural changes that affect the conformational ensemble sampled by loop 2 (**Figure 2**). Our current findings represent an atomistic mechanism behind the increased actin affinity observed in both cardiomyopathy mutations. Other cardiomyopathy mutations or myosin binding drugs may alter the conformational sampling of loop 2. For example, we reported similar allosteric effects on loop 2 for the small molecule 2’-doexy-ATP when it is in the nucleotide binding pocket of myosin, resulting in enhanced binding to actin^7,12,30,31^. Thus, loop 2 coiling may represent an important modality for understanding myosin function and we have developed methods this in the context of cardiomyopathy linked mutations that could easily be expanded to include other mutations and to study small molecules for drug development.

### Computational Predictions from Disordered Data

Simulation to simulation variability and disordered dynamics are dual features and limitations of atomic scale simulations. The intrinsic disorder of loop 2 prevents its full resolution in X-ray crystal structures and leads to high variability within and between computational simulations. While our simulations allowed for an overview of loop 2 movement and generated insights into how mutations may influence loop 2 dynamics, they do not exhaustively sample the conformational space of loop 2 (i.e. *the sampling problem*^32^). As illustrated here, even conformations of this loop that are transiently sampled can have dramatic impacts on myosin function. Less compact, more extended conformations of loop 2 are more structurally variable but may increase the probability of an association event between actin and myosin via the ‘fly casting mechanism’^33^ of intrinsically disordered protein regions. Despite the *sampling problem* our work shows a strong correlation between the orientation of loop 2 and the associative force between actin and myosin prior to weak binding. Our simulations suggest molecular mechanisms by which certain disease-associated mutations affect cross-bridge cycling. Future efforts should use longer myosin simulations or enhanced sampling strategies to better assess loop 2 conformational space. Loop 2 is also known to have a structural role in cross-bridge cycling kinetics. Simulations like those performed here should improve our understanding of the functional roles of loop 2.

While our simulations are imperfect and incomplete, they are useful to serve for generating hypotheses for future exploration. First, loop 2 may act as an electrostatic ‘tractor beam’ to guide myosin motors toward their binding sites on the thin filament. Second, the disorder of loop 2 may allow it to moonlight in several roles with different behaviors in unbound, strongly-bound, and weakly-bound conformations^16,34^. Finally, prior simulations and structures indicate that loop 2 can form interactions between myosin heads in the interacting heads motif state^2^ and it remains to be seen whether these interactions meaningfully contribute to IHM stability, and consequently thick-filament regulation. As the next generation of myosin targeting drugs are developed, their intended or unintended allosteric effects on myosin structures such as loop 2 should be considered. At the molecular level myosin targeted compounds alter myosin’s function allosterically. The present study has clearly shown that allosteric effects can propagate through myosin and alter the coiling of loop 2 to changing the affinity of myosin for actin. Drugs that directly interact with myosin have the potential to target heart disease at a mechanistic level of individual proteins, which is very exciting. As the importance of loop 2 conformational dynamics is recognized, identifying the mutations and molecules that change loop 2 kinetics through local and long-range effects is critical and the work presented here defines a paradigm for this analysis.

## Acknowledgements

This research was supported by the University of Washington Center for Translational Muscle Research (CTMR) via the National Institute of Arthritis and Musculoskeletal and Skin Diseases of the National Institutes of Health award nos. P30AR074990 and R01HL128368 to M.R. and award nos. NHLBI R01 HL141187, HL142624, and HL162229 to JD. MCC was supported by grants T32HL007828 and K99HL173646 from the National Heart, Lung, and Blood Institute. The content is solely the responsibility of the authors and does not necessarily represent the official view of the NHLBI or the NIH.

## Conflicts of Interest

The authors have no financial conflicts of interest that may be construed to these data.

## Author Contributions

Conceptualization – MCC MR

Methodology – MCC

Project Administration – MR JD

Resources – MR JD

Supervision – MCC MR JD

Visualization – KZR KJF RS

Original Draft Preparation – KZR

Review and Editing – KZR KJF MCC MR JD

Data Curation – KZR MCC

Formal Analysis – KZR MCC KJF RS

Funding Acquisition – MR JD MCC

Investigation – KZR MCC

## Methods

### Model Preparation

Homology models of pre-powerstroke (M.ADP.P_i_) human cardiac myosin (gene: MYH7) were prepared with *Modeller*^35^ using the a bovine cardiac myosin structure as a template. The template structure was the 2.45 Å X-ray crystal structure of bovine cardiac β-myosin in complex with the essential light chain, ADP, vanadate, and Mg^2+^ solved by Planelles-Herrero *et al*. (PDB: 5N69) ^36^. Crystallographic waters and other molecules were removed. Trimethylated lysine residues were converted to lysine. The X-ray study by Planelles-Herrero *et al*. included a complementary structure of pre-powerstroke myosin in the absence of OM, but residues in the tail and ELC were not resolved (PDB: 5N6A), so we used the OM-bound structure to generate our model but removed OM prior to model generation. The bovine ELC found in 5N69 was similarly used as a template for the human ELC. HIS protonation states at pH 7.0 were predicted using the *H++* webserver ^37^. Structures of the E525K and V606M myosin were generated by computationally introducing mutations into the modeled structure. This yields three separate all-atom molecular systems: WT, E525K, and V606M myosin all containing ADP.Mg^2+^.Pi in the active site and all in complex with ELC.

### Molecular Dynamics Simulation

The resulting systems were prepared for molecular dynamics simulation using the Amber 20^38^ simulation package and the ff14SB force field^39^. Water molecules were treated with the TIP3P force field^40^. Metal ions were modeled using the Li and Merz parameter set^41–43^. ADP and Pi (modeled as H_2_PO_4_)^44^ molecules were treated with the GAFF2 force field^45^. Partial charges for ADP and P_i_ were derived from a restrained electrostatic potential (*resp*) fit to quantum mechanics calculations performed with ORCA^46^. The SHAKE algorithm was used to constrain the motion of hydrogen-containing bonds^47^. Long-range electrostatic interactions were calculated using the particle mesh Ewald (PME) method. Hydrogen atoms were modeled onto the initial structure using the *leap* module of *AMBER,* and each protein was solvated with explicit water molecules in a truncated octahedral box and neutralizing counterions were added. Each system was minimized in three stages. First, hydrogen atoms were minimized for 1000 steps in the presence of 100 kcal/mol restraints on all heavy atoms. Second, all solvent atoms were minimized for 1,000 steps in the presence of 25 kcal/mol restraints on all protein atoms. Third, all atoms were minimized for 8,000 steps in the presence of 25 kcal/mol restraints on all backbone heavy atoms (N, O, C, and C_α_ atoms). After minimization, systems were heated to 310 K using the NVT (constant number of particles, volume, and temperature) ensemble and in the presence of 25 kcal/mol restraints on backbone heavy atoms. Next, the systems were equilibrated over five successive stages using the NPT (constant number of particles, pressure, and temperature) ensemble. During the first four stages, the systems were equilibrated for 5 ns in the presence of 25 to 1 kcal/mol restraints on backbone heavy atoms. During the final equilibration stage, the systems were equilibrated for 5 ns in the absence of restraints. A 10 Å nonbonded cutoff was used for all preparation and production simulations. Separate equilibrations were run for replicate simulation for each genotype. The equilibrated systems were then simulated using conventional molecular dynamics protocols in the NVT ensemble in triplicate for 500 ns each (3 systems, three replicate simulations per system, 500 ns sampling per replicate simulation = 4.5 µs total sampling) and coordinates were saved every 10 ps.

### *DelPhiForce* Calculations

To assess the impact of loop 2 residue variations on electrostatic interactions, we employed *DelPhiForce*^8,48^ to compute the electrostatic binding forces between representative timeframes extracted from MD simulations and a pre-equilibrated actin pentamer derived from the 8EFH PDB structure^24^. *DelPhiForce* calculates electrostatic forces by numerically solving the Poisson-Boltzmann (PB) equation, which accounts for the distribution of ions and the dielectric properties of the solvent and protein. This approach allowed us to determine the electrostatic contributions to the interaction energy between the loop 2 variants and the actin pentamer, providing insights into how residue substitutions modulate binding affinity.

Alignment of myosin with actin in a pre-weakly bound sate was achieved using a reference structure of the human cardiac actin-tropomyosin-myosin complex in complex with ADP-Mg2+ (PDB: 8EFH). All alignments were generated using the MatchMaker function in Chimera (UCSF). An equilibrated actin pentamer was aligned to actin such that the central actin monomer was interacting with myosin. Frames from each simulation (with a resolution of 1 frame/ns) were then aligned to actin using the 8EFH reference. Importantly the reference structure represents a strongly bound actomyosin complex. To generate a weakly bound conformation myosin structures from MD simulation were only aligned to the HLH motif of the reference structure (residues 515-550 and 530-553 respective). Myosin was then shifted 30 Å away from action to simulate pre-binding conditions. To achieve this a plane of actin and myosin interaction was defined by the list of clashing atoms between the two molecules. Myosin was then shifted 30 Å normal to this plane away from actin. This alignment can be seen in **Supplemental Figure 3**. Protein structures were prepared for electrostatic modeling using *DelPhiPka* ^49^ and *DelphiForce* was used to calculate electrostatic binding force. Default parameters were used for all *Delphi* programs.

### Molecular Dynamics Analyses

Dihedral angles were calculated for every residue in loop 2 in every timestep of every simulation using the *cpptraj* command *multidihedral* for residues 615-651. Based on resulting Ramachandran coverage loop 2 was defined as residues 621-646 (**Supplemental Figure 2**). The center of mass of the positively charged Lys residues in loop 2 was calculated using the *cpptraj* command *vector* to identify the center of mass for the C_α_ of residues 633, 635, 637, and 639 (**Figure 4A**). The location of the center of mass of the negatively charged N-terminal residues in the actin molecule used was calculated using the *define centroid* command in *Chimera* based on the C_α_ of residues 2-5. Finally, *cpptraj* and the *contacts* command was used to calculate the contact pairs for each residue in myosin simulations. Two residues were considered in contact with one another if at least one pair of heavy atoms were within 5 Å of one another.

### Principal Component Analysis of Loop 2 C_α_ Position and Loop 2 Dihedral Angles

Principal component analysis was used to define loop 2 position and coiling as previously described^50–52^. Briefly, to account for movement of the myosin molecules during simulation, all myosin molecules used for analysis (1 frame/ns from each simulation) were aligned to their 50 kDa domains (residues 215-231, 266-453, 604-621, 645-665, 472-590). The C_α_ position of each residue as well as the dihedral angels of each residue in loop 2 was then extracted from the simulations (residues 621-646). These data were then used for PCA analysis. For the dihedral angle PCA the first 100 ns (100 frames) of each simulation were omitted to allow the simulations to acclimate as they were highly variable. Additionally, the four glycine residues in loop 2 were omitted (residues 626, 634, 636, 641) since glycine dihedral angels are extremely variable. Additionally, the dihedral angle PCA was performed on the sin() and cos() components of the phi/psi dihedral angles to avoid circular statistics as previously described^53^. All frames/timesteps and all loop 2 residues were used for the Cartesian coordinate C_α_ PCA analysis.

## Supplemental Figures

**Supplemental Figure 1:**
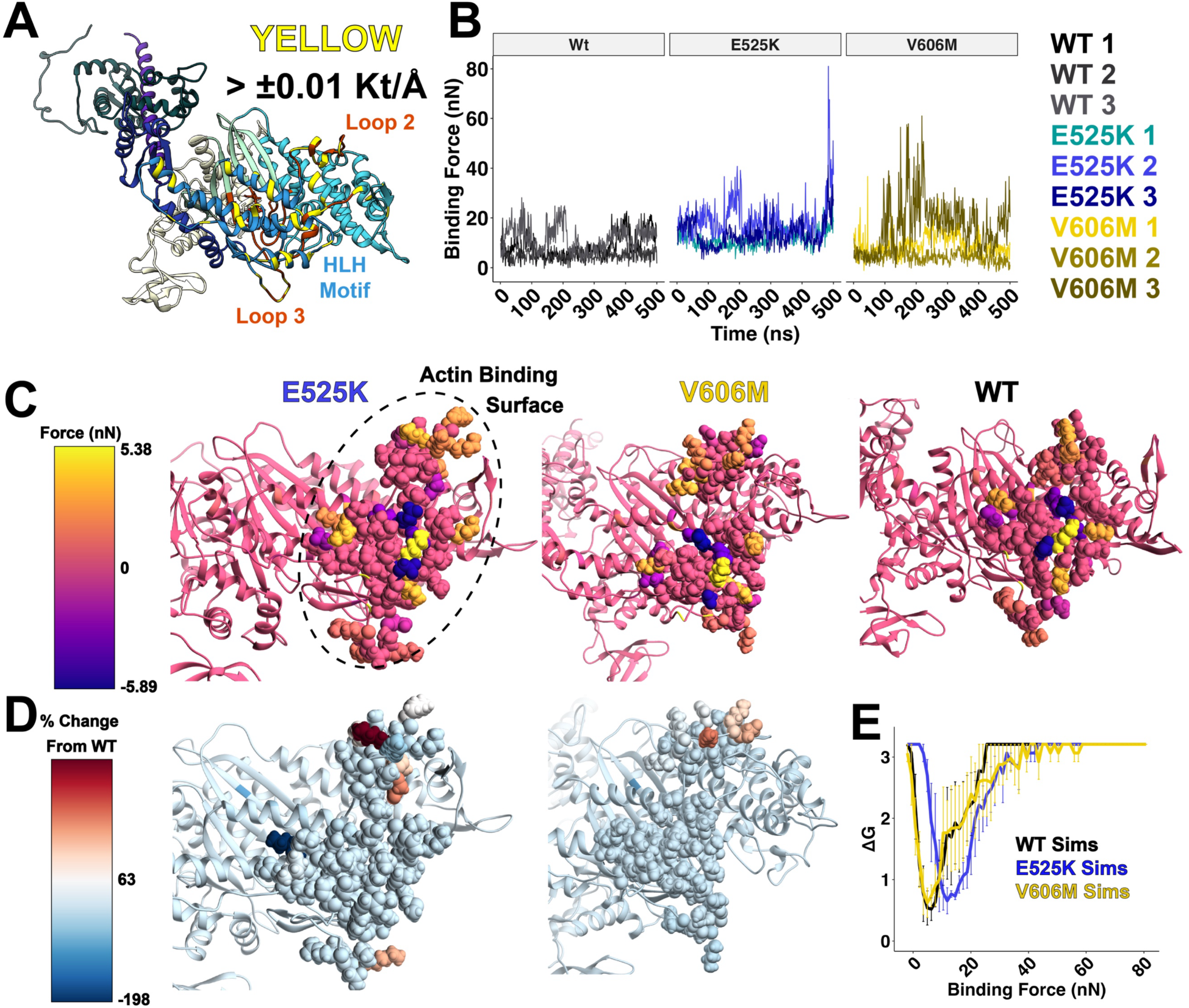
Binding force felt by each residue in the actin binding surface. (A) Pre-powerstroke human β-myosin S1 head where residues that experienced > ±0.0 kT/Å during any (WT or Mutant) *DelPhiForce* simulation are highlighted in yellow. Key charged residues in the actin binding surface can be seen including the HLH motif, loop 2, loop 3, and the strut. (B) Binding force calculated with DelphiForce across simulation time in each of three replicate simulations of each genotype. (C) Myosin S1 actin binding surface colored by average binding energy felt at that residue during three replicate simulations. Loop 2, loop 3, and the HLH motif are depicted with a space filling model to highlight the changes in these areas. (D) Myosin S1 actin binding surface depicted as in C but colored by percent change in average binding force felt by that residue when compared to WT simulations. (E) Boltzmann free energy distribution showing the distribution of WT, E525K, and V606M simulations that sample different binding energies for the whole myosin S1 head.

**Supplemental Figure 2:**
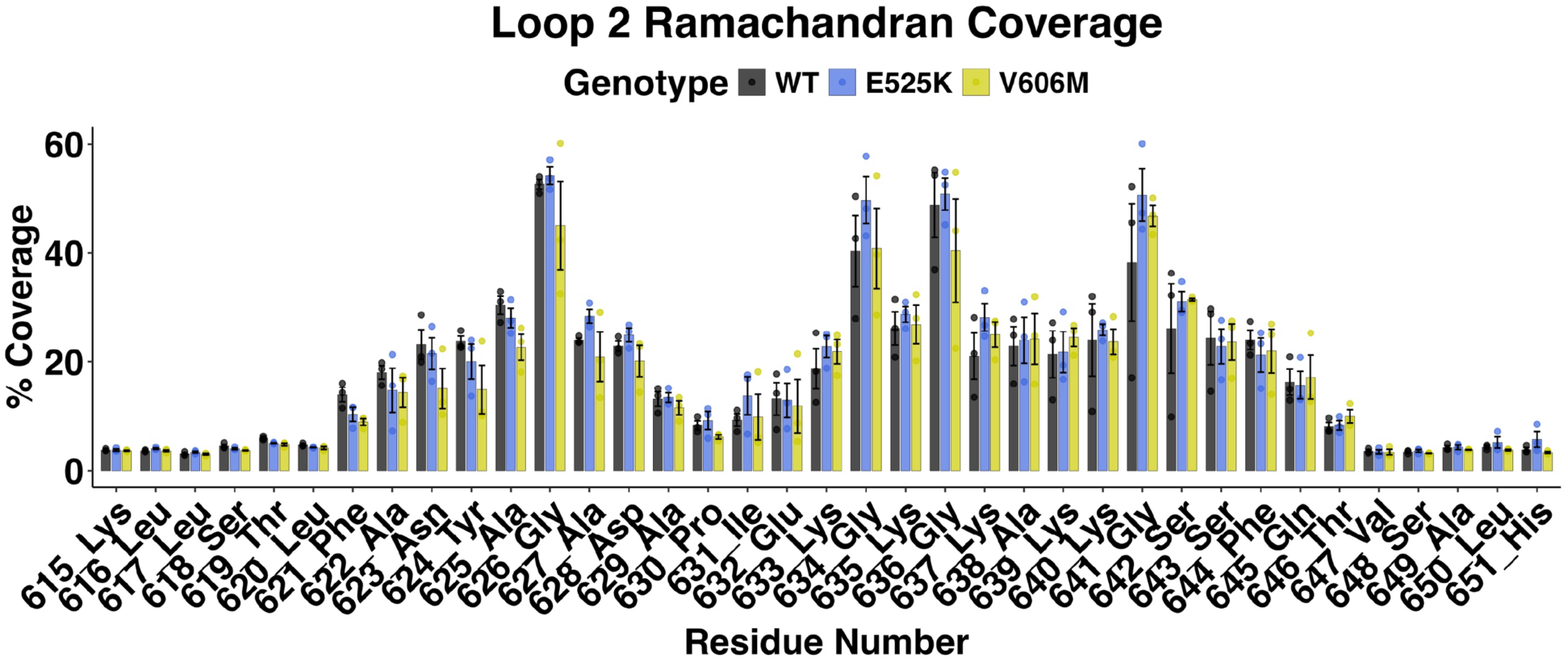
Identification of loop 2 residues in simulation. Dihedral angles were recorded for resides in the vicinity of loop 2 for each simulation. Loop 2 was defined for all myosin molecules in all simulations as residues 621 to 646 due to the increase in Ramachandran plot coverage in this region.

**Supplemental Figure 3:**
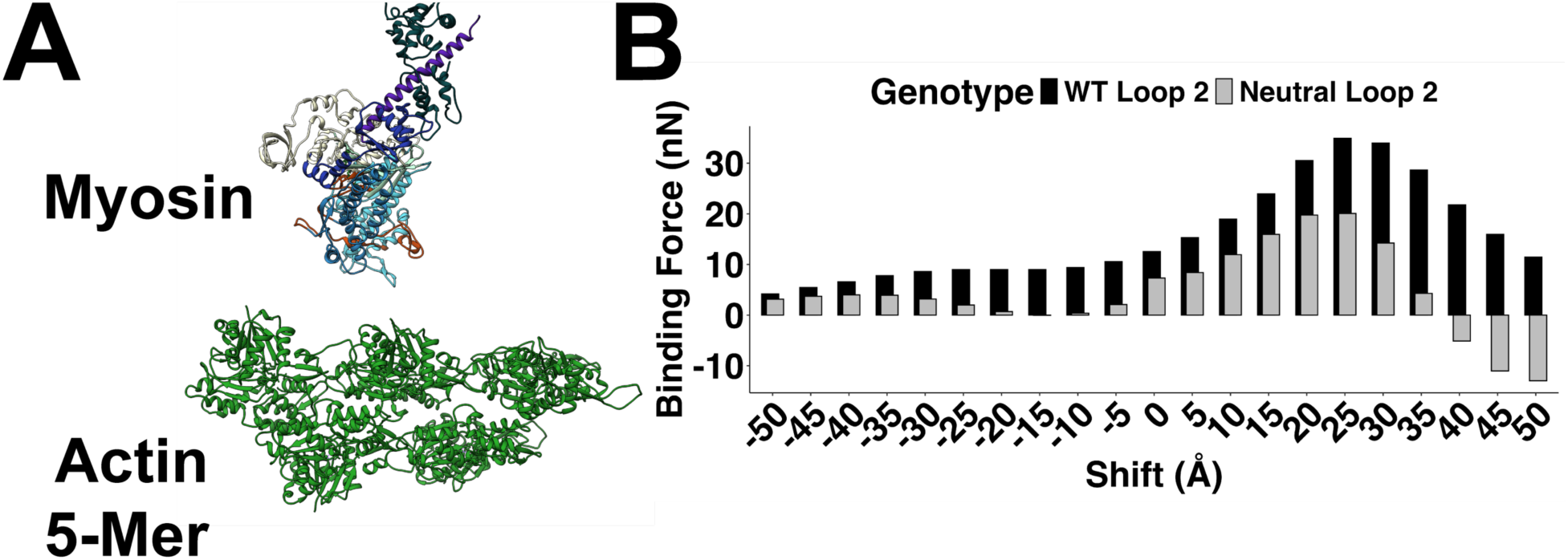
Setup of *DelPhiForce* simulations and validation of *DelPhiForce*. The alignment of myosin and actin was achieved as discussed in the methods section. (A) A depiction of the alignment of a WT myosin S1 head 30 Å from its binding site on an actin 5-mer. (B) To confirm that the *DelPhiForce* tool would model electrostatic interactions between myosin and actin a test simulation was run where a WT myosin S1 head was shifted along the axis of actin and the binding force was calculated after each 5 Å shift (black in B). Additionally, to confirm this tool could identify the electrostatic force generated by the charged lysine residues of loop 2 these residues were computationally changed to glutamine residues (630, 632, 634, 636, and 637). A reduction in the binding force between myosin and actin can be seen when the lysine residues of loop 2 are neutralized (gray in B).

**Supplemental Figure 4:**
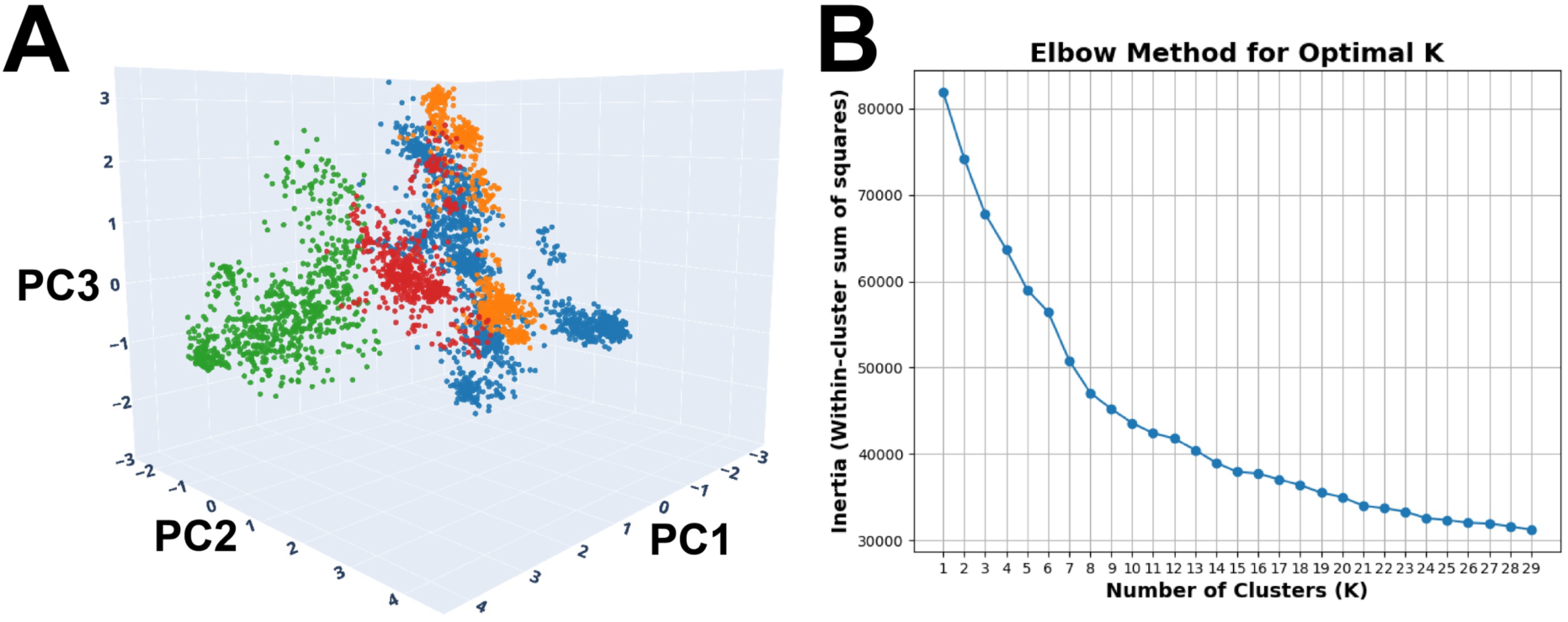
K-means clustering of loop 2 dihedral angle PCA and loop 2 secondary structure. 3D depiction of the four clusters identified showing distinct grouping of clusters. (B) Inertia plot (within-cluster sum of squares) used to determine optimal number of clusters for k-means analysis.

## Notes

### Competing Interest Statement

The authors have declared no competing interest.

